# Large-scale discovery of mouse transgenic integration sites reveals frequent structural variation and insertional mutagenesis

**DOI:** 10.1101/236307

**Authors:** Leslie O. Goodwin, Erik Splinter, Tiffany L. Davis, Rachel Urban, Hao He, Robert E. Braun, Elissa J. Chesler, Vivek Kumar, Max van Min, Juliet Ndukum, Vivek M. Philip, Laura G. Reinholdt, Karen Svenson, Jacqueline K. White, Michael Sasner, Cathleen Lutz, Stephen A. Murray

## Abstract

Transgenesis has been a mainstay of mouse genetics for over 30 years, providing numerous models of human disease and critical genetic tools in widespread use today. Generated through the random integration of DNA fragments into the host genome, transgenesis can lead to insertional mutagenesis if a coding gene or essential element is disrupted, and there is evidence that larger scale structural variation can accompany the integration. The insertion sites of only a tiny fraction of the thousands of transgenic lines in existence have been discovered and reported due in part to limitations in the discovery tools. Targeted Locus Amplification (TLA) provides a robust and efficient means to identify both the insertion site and content of transgenes through deep sequencing of genomic loci linked to specific known transgene cassettes. Here, we report the first large-scale analysis of transgene insertion sites from 40 highly used transgenic mouse lines. We show that the transgenes disrupt the coding sequence of endogenous genes in half of the lines, frequently involving large deletions and/or structural variations at the insertion site. Furthermore, we identify a number of unexpected sequences in some of the transgenes, including undocumented cassettes and contaminating DNA fragments. We demonstrate that these transgene insertions can have phenotypic consequences, which could confound certain experiments, emphasizing the need for careful attention to control strategies. Together, these data show that transgenic alleles display a high rate of potentially confounding genetic events, and highlight the need for careful characterization of each line to assure interpretable and reproducible experiments.

## INTRODUCTION

Since the report of the production of the first germline competent transgenic mouse more than 35 years ago(Gordon and Ruddle 1981), transgenic mouse models have had an enormous impact on biomedical research, providing a range of tools from critical disease models to more broadly useful reporters and recombinase-expressing lines. The majority of transgenic lines are produced through microinjection of the desired DNA fragment into the pronucleus of a zygote, although lentiviral transgenesis and production through an ES cell intermediate has been reported and used to some extent(Pease et al. 2011). Typically, transgenes comprise engineered DNA fragments ranging in size from small plasmid-based constructs to much larger bacterial artificial chromosomes (BACs), which insert into the genome in a presumably random fashion, usually as a multi-copy array. Founder lines are then examined for both transmission and for the desired expression levels and specificity, often leading to the rejection of many lines that fail to express the transgene properly. While the mechanism for this variation in outcome is unclear, it is presumed that genetic context of the integration locus plays some role in providing a transcriptionally permissive environment. There are many additional factors that could affect transgene expression, including copy number, and thus ultimately selection of founders is an empirical exercise and often only a single line is chosen for experiments and publication.

Of the 8,012 transgenic alleles published in the Mouse Genome Database, only 416 (5.2%) have an annotated chromosomal location. For transgenic *cre* alleles, the number is even lower, with a known chromosomal location for 36/1,631 (2.3%) lines, highlighting the challenge of identification of integration sites despite widespread acknowledgement that such information is useful and important. Low resolution mapping of transgenes can be achieved through FISH or linkage mapping, but these approaches offer little information about potential mutagenesis at the integration site. Inverse PCR can be used to clone the actual fusion sequence, but has a high failure rate owing to the multi-copy nature of most transgenes. More recently, high-throughput sequencing (HTS) has been employed to identify transgene insertion sites (Dubose et al. 2013), with improvements offered by the use of mate pair libraries(Srivastava et al. 2014). Despite the promise, HTS-based approaches have not seen widespread implementation, possibly due to the cost and/or complexity of the analysis.

The identification of transgene insertion sites is useful for a number of reasons. First, it allows the user to avoid experimental designs that attempt to combine linked alleles (e.g. a conditional allele with a cre transgene), obviating a long and possibly fruitless breeding exercise. Second, it enables the design of allele-specific genotyping assays, which assist in colony management and determination of zygosity. Finally, it alerts the investigator to potential confounding effects of insertional mutagenesis through the direct disruption of the coding sequence of endogenous genes, indirect effects on the regulation of nearby genes, or complex structural variations (inversions or duplications) that can accompany the integration event. Cases of insertional mutagenesis with dramatic phenotypic consequences have been reported. For example, the Tg(TFAP2A-cre)1Will allele inserted into the *Hhat* gene, disrupting its function, leading to a variety of severe developmental abnormalities in homozygous embryos including holoprosencephaly with acrania and agnathia, reflecting a disruption of the hedgehog signaling pathway (Dennis et al. 2012). Given the utility of this line in targeting branchial arches of the developing face, this could confound the interpretation of experimental data if the correct breeding scheme and controls are not included. Because so few insertion sites have been mapped, the scale of this issue is unknown. A prior report using FISH found that transgenes tend to insert into G-positve band regions (Nakanishi et al. 2002), which typically have reduced gene density, but the mapped transgenes were not assessed for expression levels, so it is unclear if these data are representative of transgenes used in the wider scientific community. More recently, Targeted Locus Amplification (TLA) (de Vree et al. 2014; Hottentot et al. 2017) has been employed to identify the insertion site for 7 Cre driver lines(Cain-Hom et al. 2017), only one of which was found to insert into an annotated gene. However, because of the small sample size, it is not clear if this rate of mutagenesis is representative of the genome-wide rates in larger collections representing a variety of transgene types.

Here, we describe the first large-scale survey of insertion sites in widely used transgenic mouse lines. The TLA capture process is highly efficient, providing high-resolution localization of the transgene insertion site for all 40 lines tested, including 24 with fusion reads discovered on both ends of the insertion. Remarkably, half of the insertion events disrupt one or more genes. Some of these disrupted genes have known knockout phenotypes, including embryonic lethality. We also identify a number of structural variations, including frequent deletions and duplications, and unexpected elements that have cointegrated with some of the transgenes. Finally, we show that transgenic lines can display phenotypes independent of experimental context, consistent with our insertional mutagenesis discoveries. Together, these data demonstrate the clear need to consider the molecular consequences of transgene insertion in any experimental design.

## Results

We selected a total of 40 transgenic lines from live colonies in the JAX Repository for our study, including 4 lines distributed through the Mouse Mutant Research and Resource Center (MMRRC) at JAX. All lines are broadly utilized and thus represent important research tools that would benefit from insertion site identification. The list comprises 17 genetic tool strains, including 15 cre drivers, many of which have demonstrated off-target or unexpected excision activity (Heffner et al. 2012). In addition, we included 5 lines that lack an allele-specific genotyping assay, and 18 critical Alzheimer’s or Parkinson’s disease models (Figure 1A). We selected lines that were generated through a variety of means (Figure 1B), including standard small plasmid-based transgenes, human and mouse BAC transgenes, a human PAC, a human cosmid, and a transgene generated through lentiviral-mediated transgenesis. The BAC/PAC/cosmid vectors were included both to capture critical lines of interest and to test the feasibility of the TLA process on these larger constructs. A schematic of the TLA process, depicted in Figure 1C, was performed essentially as described ((de Vree et al. 2014; Hottentot et al. 2017); Materials and Methods) using primers specific for known elements of each transgenic line (Supplementary Table 1). Transgene insertion sites result in high sequencing coverage across the transgene and its insertion site(s), and at least one putative fusion read across the transgene-genome breakpoint was identified for all 40 lines (Table 1; Supplementary Table 1). In some cases, follow-up TLA analysis using additional primers designed based on the initial results was required to identify a fusion read at either junction. Insertions were found genome-wide on 17/19 autosomes, with 5 insertions each identified on chromosomes 1 and 2 (Figure 1D). Structural variations accompanying the insertion were identified for a majority of lines (30/40), comprising 24 deletions and 6 duplications (Figure 1E). As some of the fusion contigs were constructed using single reads, we used PCR and Sanger sequencing to verify fusion reads identified by TLA. Overall, we identified and confirmed both fusion breakpoints for 19/40 transgenes, and a single fusion breakpoint for an additional 21, demonstrating the efficiency of the TLA process in identifying the precise insertion sites (Supplementary Table 1 & 2). For deletions where only one fusion read could be confirmed, a quantitative PCR loss of native allele (LOA) (Valenzuela et al. 2003; Frendewey et al. 2010)assay was used to confirm the loss of either the genes within the deletion (see below) or a region close to the estimated insertion site (Supplemental Table 1 & 3). Remarkably, the transgene insertion event in 20 of 40 lines either delete at least one exon of one or more genes (14) or insert into an intron likely affecting its normal transcription (6). Overall, these data indicate a strong enrichment of transgene insertion events in genic regions of the genome, placing these lines at high risk for confounding phenotypes due to insertional mutagenesis.

**Figure 1.**
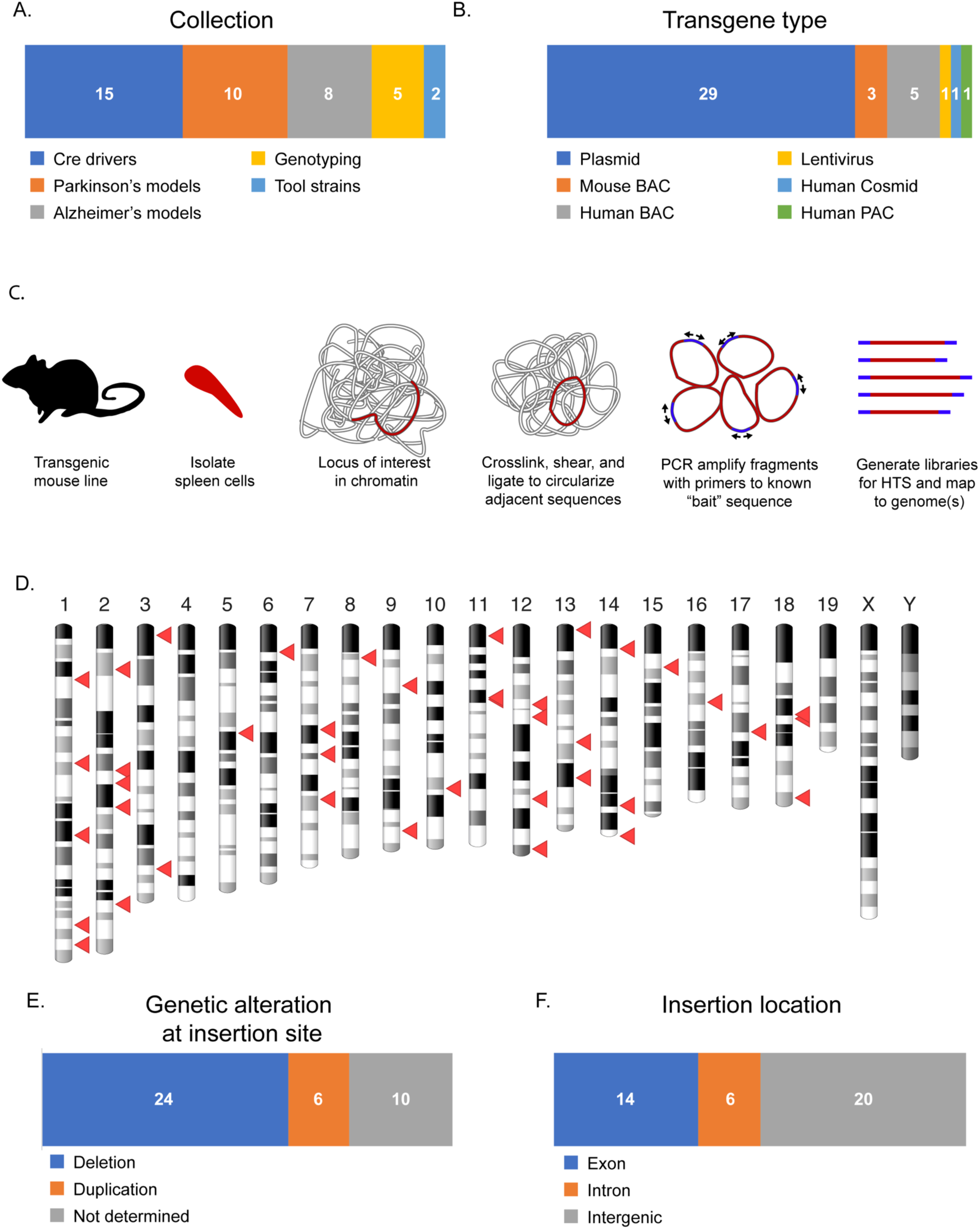
Discovery of the integration loci for 40 transgenic mouse lines. A. Distribution of the categories of transgenes included in this study. B. Distribution of transgenes by molecular type. C. Schematic of the targeted locus amplification process. C. Ideogram showing the physical distribution of transgene insertion sites identified by TLA. D. Types of genetic alterations that accompany transgene insertions. E. Proportion of insertion sites that occur in genes (exon or intron) or non-gene loci (intergenic).

**Table 1.**
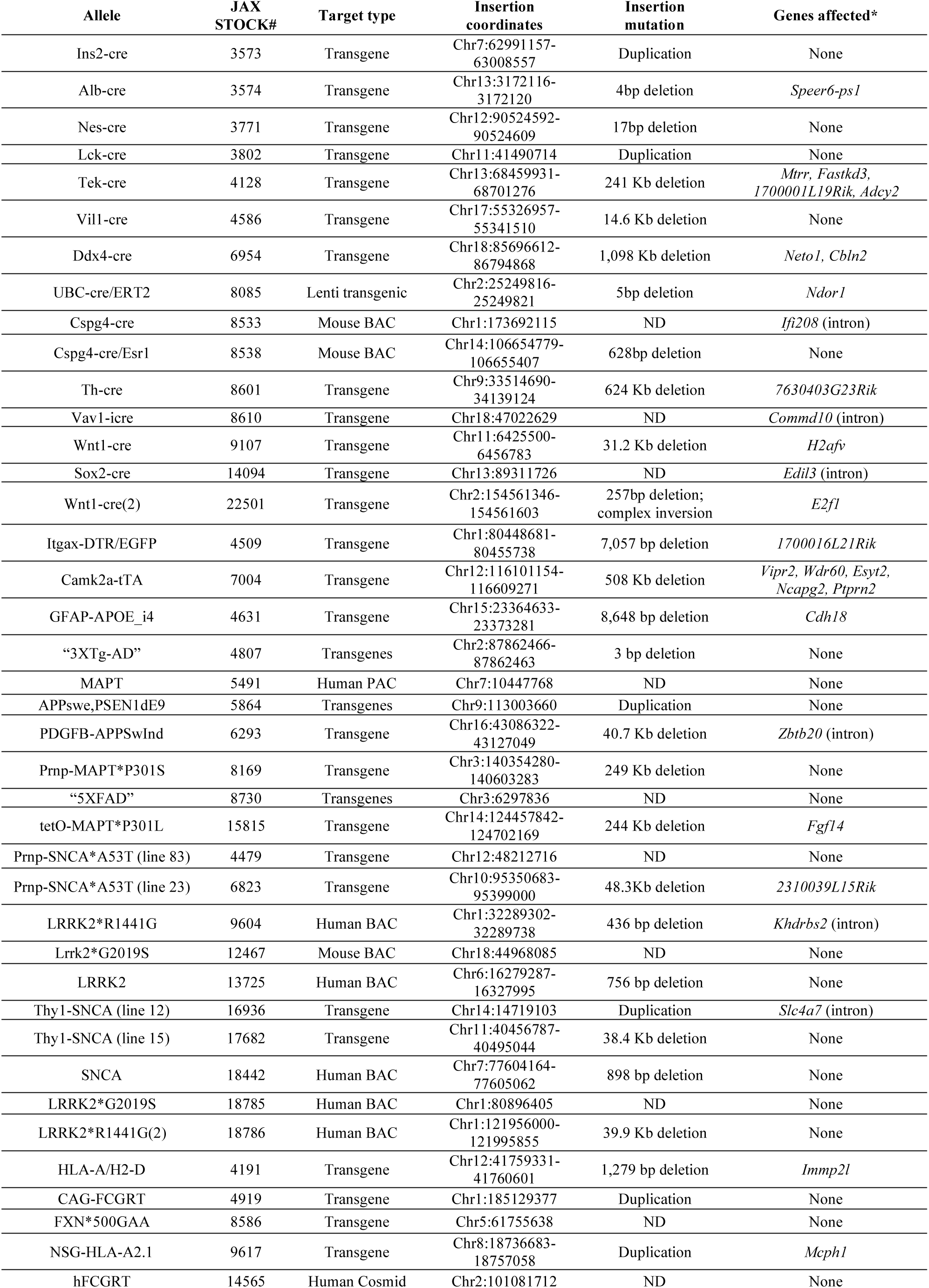
Summary of transgene insertion sites identified in this study

A majority of insertion events discovered were accompanied by a deletion (Figure 1E), which varied in size from a few basepairs to 1.1Mb in the case of the Ddx4-cre line (Figure 2A). As noted above, amongst the 24 deletions, we identified a surprisingly high rate of insertional mutagenesis, either deleting or disrupting between 1 and 5 mouse genes (Figure 2B), each with potential phenotypic consequences. In addition, two genes are disrupted through duplication event that accompanied their respective insertion (Supplementary Table 1). Of the total 28 genes disrupted, 15 have a previously reported knockout (KO) phenotype, including 4 with embryonic lethal phenotypes (Figure 2C), and multiple genes are deleted in 3 lines, highlighting the potentially confounding effect of the insertion event. For example, in the Tek-cre line we identified a 241Kb deletion on Chromosome 13 that includes four protein coding genes (*Mtrr*, *Fastkd3*, *1700001L19Rik*, *Adcy2*) (Figure 2 D,E). We validated both breakpoints and confirmed the deletion of *Mtrr*, *Fastkd3*, and *Adcy2* by loss-of-native-allele qPCR assays, showing clear loss of one copy for all three genes, and in addition confirming the breakpoint between exons 14 and 15 in *Adcy2* (Figure 2F). In addition to the frequent widespread (off-target) activity seen in this line (Heffner et al. 2012), prior reports show that an *Mtrr* gene trap allele exhibits trangenerational epigenetic effects leading to severe developmental abnormalities when breeding from a female carrier (Padmanabhan et al. 2013). Given the common use of this line to analyze vascular development, the transgene itself could confound analysis depending on the breeding scheme, highlighting the need for proper controls (i.e. Cre-only) in studies using this transgene. Finally, we confirmed the deletion of an additional 13 genes (Figure 2G, Supplementary Table 1), including two (*Immp2l* and *Cdh18*) that appear null due to the maintenance of the line as a homozygote. Interestingly, one line (PDGFB-APPSwind) showed insertion into the gene *Zbtb20*, but a recent report clearly shows that despite the insertion of more than 10 copies of the transgene, expression of the protein in heterozygous transgenic mice is comparable to WT, suggesting other regulatory mechanisms to maintain a uniform level of expression (Tosh et al. 2017). Therefore, in some cases the consequences of intron insertion require independent validation. Together, these data show that transgene insertions are often associated with large mutagenic deletions, affecting one or more genes, potentially confounding interpretation of results unless the proper control strategies are employed.

**Figure 2.**
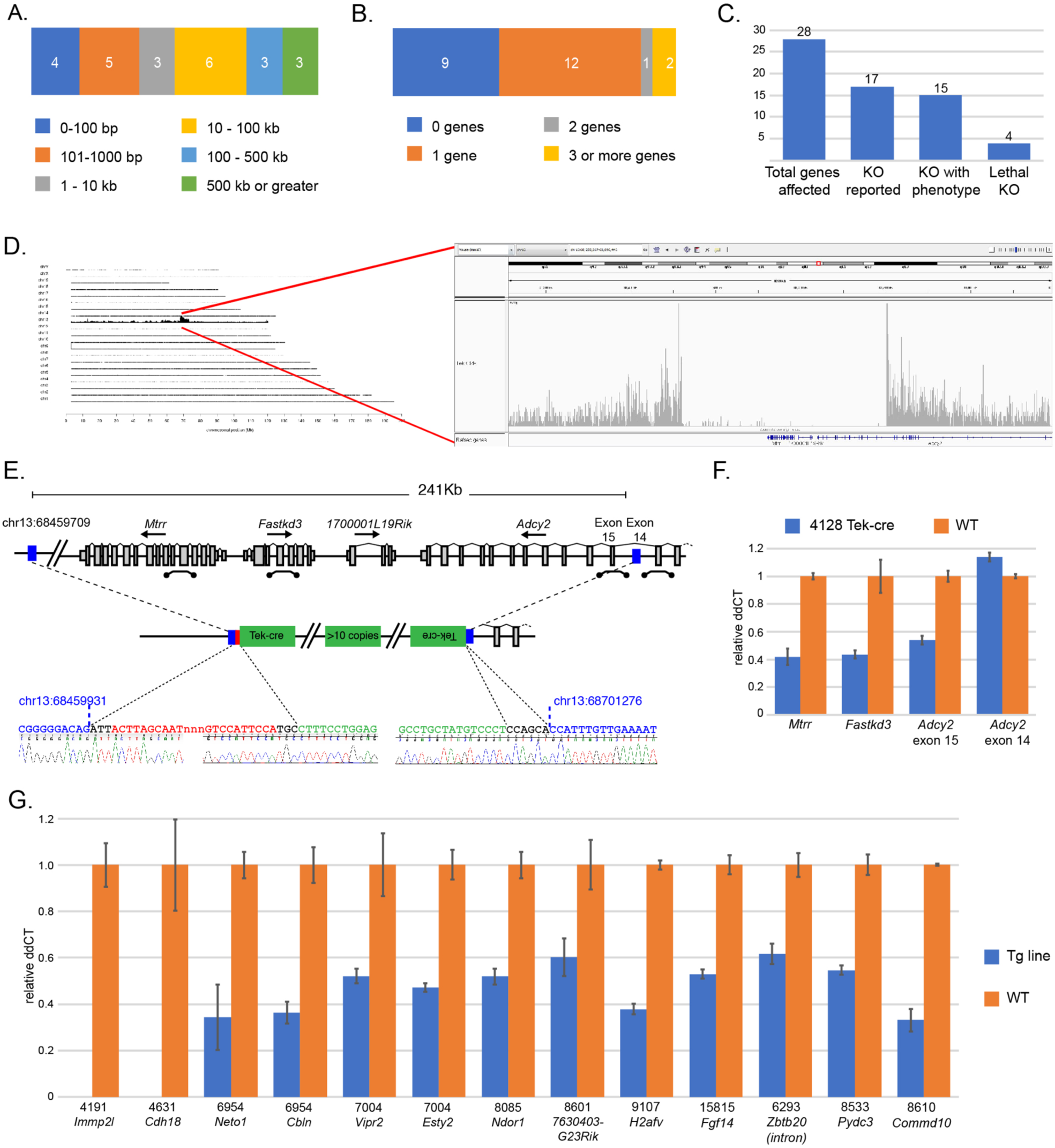
Deletions accompanying transgenic insertions. A. Profile of sizes of deletions identified at integration loci. B. For integrations that occur in genes, the profile of the number of genes affected by the insertion event. C. Illustration of the potential impact of transgene insertions into genes, including the number of genes with reported knockout (KO) alleles, the number of KO alleles with a reported phenotype, and number of genes shown to be essential for life. D. Schematic of the insertion locus in the Tek-cre line. Blue bars indicate the 5’ and 3’ limits of the deleted region, with the relative orientation of transgene copies adjacent to the breakpoint as determined from sequence-confirmed fusion reads. Locations of qPCR probes to confirm copy number are shown. E. Results of loss of native allele (LOA) qPCR assays showing the expected loss of one copy of *Mtrr* and *Fastkd3* and exon 15 of *Adcy2*, which lie within the deletion. *Adcy2* exon 14, which lies outside of the deletion, has the expected two copies. WT copy number is arbitrarily set at 1, thus a value of 0.5 would indicate loss of one copy. E. LOA assays for 13 other genes/loci deletions identified in this study. Strains are indicated by Stock # above the gene symbol for each test. For strains 4191 and 4631, the complete loss of *Immp2l* and *Cdh18*, respectively, is consistent with the homozygous maintenance of these lines.

TLA analysis also revealed additional structural variations around the insertion site, including 6 instances of duplications and one inversion accompanied by a large deletion. In many duplication cases, fusion reads were only identified on one end of the transgene insertion, so the exact extent of the duplication could only be estimated through read depth. However, we were able to confirm additional copies of parts of the genes *Mcph1* and *Slc4a7* (data not shown), although it is not clear how this might affect gene function. For the Wnt1-Cre(2) line(Lewis et al. 2013), we observed a complex structural variation on chromosome 2, involving a large 45Kb inverted segment inserted into exon 5 of the *E2f1* gene (Figure 3A). The inversion itself contains all of exon 5, but deletes 23Kb including exons 6 and 7 of *E2f1* proximal to the transgene insertion location, all of *Necab3*, *1700003F1Rik*, and a portion of the 3’

**Figure 3.**
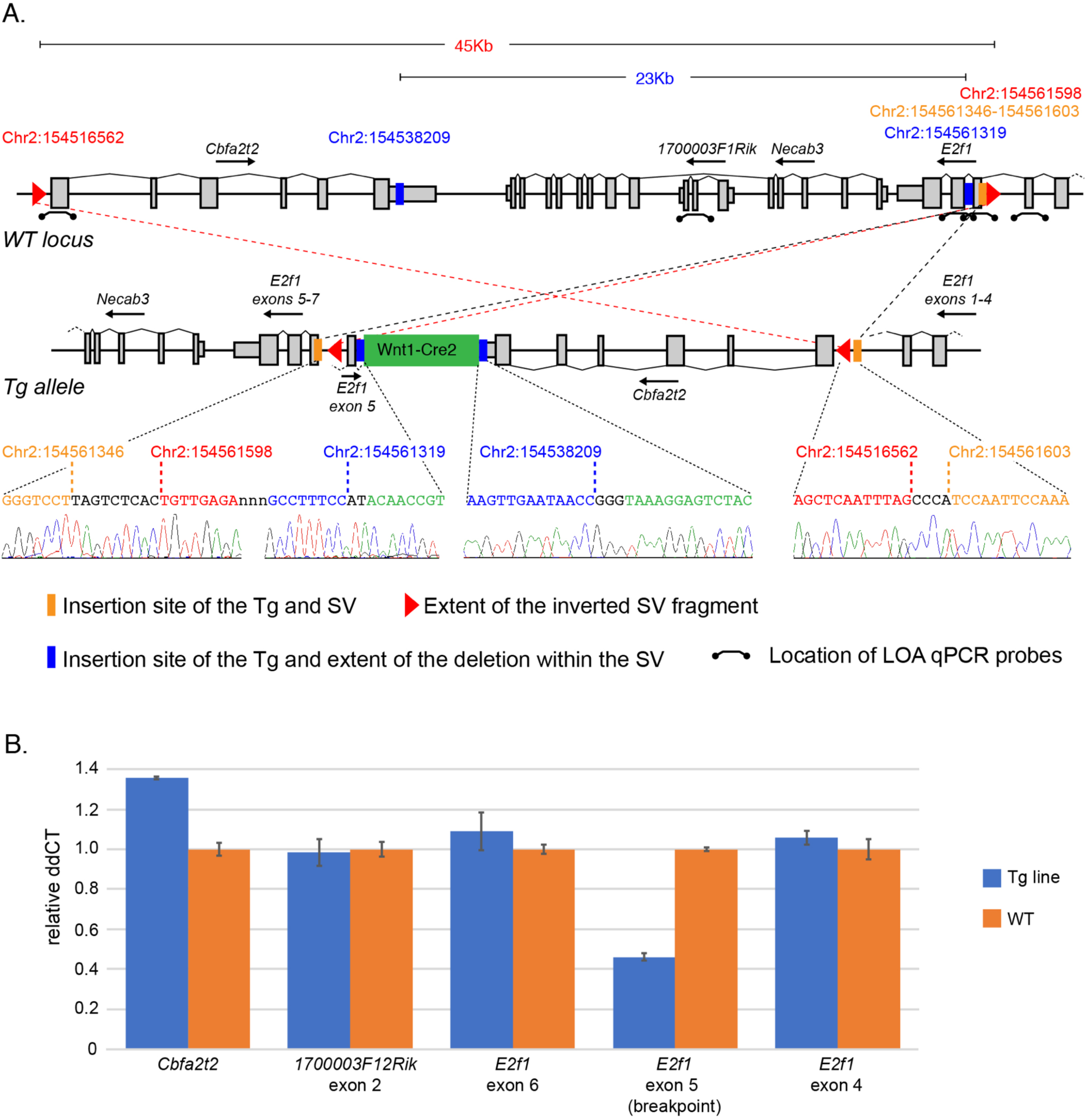
Complex structural variations accompanying transgenic insertions. A. Schematic of the structural variation accompanying the Wnt1-cre2 transgene insertion. The locus includes a large duplication with a partial deletion that accompanies the transgene insertion. The entire duplicated interval is inverted and is inserted into exon 5 of the E2f1 gene. The red triangles identify the extent of the entire SV that is inverted, the blue bars indicate the insertion site of the transgene and the extent of the deletion within the duplicated fragment, and the orange bars indicate the location of the SV insertion. qPCR probes are indicated on the WT locus. The qPCR probe for *E2f1* exon 5 spans the breakpoint of SV insertion. Confirmation of each fusion read that defines the SV by PCR-Sanger sequence is illustrated. B. LOA confirmation of the expected copy number for each gene/exon affected by the SV.

UTR of *Cbfa2t2*. As a result, the structural variation disrupts the *E2f1* gene, with the concomitant duplication of exon 5 in the opposite orientation. We used an LOA assay to confirm the disruption of exon 5 (Figure 3B), but does not capture the duplication of the inverted exon, as the inverted fragment is smaller than the amplicon of the qPCR probe. The copy number of *E2f1* exons 4 and 6 are unaffected, as they surround the structural variation. In addition, an LOA assay shows the duplication of exon 6 of *Cbfa2t2*, and an LOA for exon 2 of *1700003F12Rik*, which resides in the deleted portion of the duplicated fragment, shows the normal two copies as expected (Figure 3B). Together, these data illustrate the potential complex structural variations that can occur with transgene integration.

Because TLA isolates all DNA fragments in close proximity to the transgene integration site, it is possible to identify components of the transgene itself, in addition to the surrounding mouse sequence. The only limitation is the selection of reference genomes for mapping. In this study, we typically mapped to genomes predicted to be part of the transgene, based on the published description of the transgene construction. While for the most part we were able to identify construct elements described in the original publications, unexpected components were seen in several transgenic lines. For example, we found that an entire hGH minigene, described as a polyA sequence, was present in four lines (Ins2-cre, Alb-cre, Nes-cre, and Lck-cre; Figure 4A), as previously reported for the Nes-cre line (Declercq et al. 2015). Of note, publications for these lines reference a vector originally described in (Orban et al. 1992), which clearly describes the minigene structure of the cassette. Similarly, TLA and PCR validation of the Vil-cre line reveals the presence of the entire *Mt1* gene sequence (Figure 4B), despite its description as a “metallothionein poly(A) signal” in the original publication (Madison et al. 2002). The source plasmid does indeed describe it as containing the polyA and several introns (Sauer and Henderson 1990).

**Figure 4.**
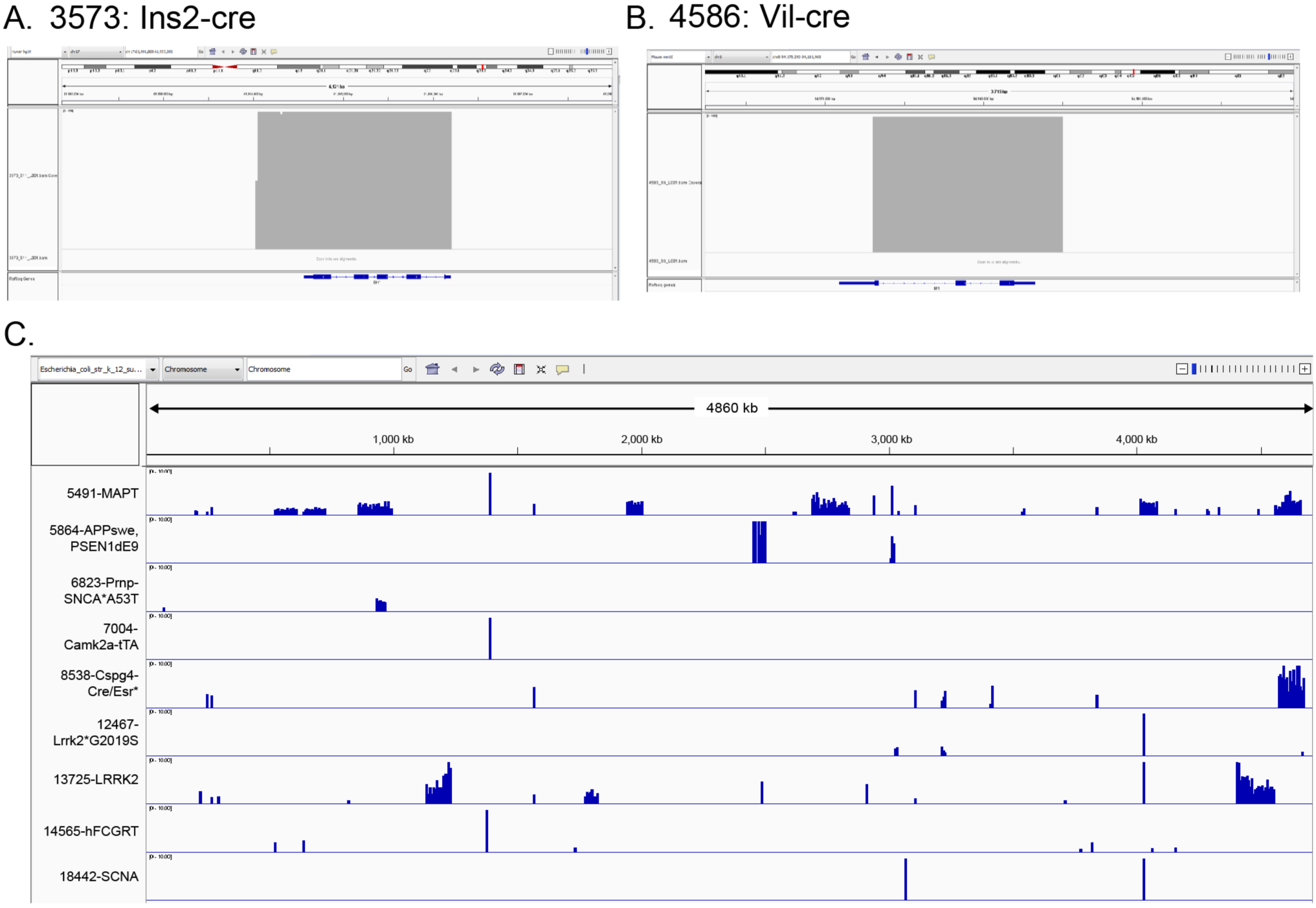
TLA reveals additional passenger cassettes and fragments in transgenes. A, B. View of TLA reads that map to the human growth hormone (hGH) gene for two transgenes (Ins2-cre and Vil-cre), showing the inclusion of the entire gene structure, including coding exons. C. Reads for nine transgenes mapped to the *E.coli* genome indicating a variable level of co-insertion into the transgene integration site. Deep coverage for discrete loci shared between multiple lines suggests that these sequences are part of the transgene vector. The amount of *E.coli* co-integration ranges from a few hundred bp to more than 200Kb.

Although the impact of the presence of the *Mt1* minigene is unclear, there is evidence that the Nes-cre hGH minigene is expressed and that this expression is responsible for some of the metabolic phenotypes observed in the Nes-cre line (Galichet et al. 2010; Giusti et al. 2014). These data indicate that TLA has the added potential to expand on and confirm reported transgene composition, and in some cases can correct, clarify, and/or update the record for these strains.

In our initial analysis, we identified several cases where mapped sequences were fused to unknown sequences. Further analysis revealed that some of these fusions were with the *E.coli* genome, and not vector sequences, suggesting that fragments of contaminating *E.coli* DNA were co-integrating with the transgene. To assess the frequency of this phenomenon, we mapped all of the TLA data to the *E.coli* genome (K-12) and found evidence for co-integration in 10/40 strains, with total composition ranging from as little as 300bp to more than 200Kb (Figure 4C). Some of the small fragments were identical between samples, suggesting that these were intentional components of the transgene, possibly as part of the vector backbone. However, for lines with significant *E.coli* genome contribution, it is likely that this is the result of contamination in the microinjection preparation of the construct. While the impact of this finding for these specific lines is unclear, prior reports have shown that bacterial sequences can contribute to transgene silencing (Scrable and Stambrook 1997; Chen et al. 2004).

While some of the small number of transgene insertion sites currently known were discovered following the serendipitous identification of an unexpected transgene-specific phenotype, systematic phenotyping of transgenic lines to assess the impact of transgene insertion has not been reported. Taking advantage of the high-throughput KOMP2 Phenotyping platform at JAX (White et al. 2013; de Angelis et al. 2015; Dickinson et al. 2016; Karp et al. 2017; Meehan et al. 2017), we asked whether we could detect phenotypes in a selection of 7 Cre driver lines from the lines examined above. As colony maintenance strategy differed amongst lines, we pooled control mice and WT C57BL/6J to create a reference population. As shown in Figure 5, we identified 66 significant phenotypes amongst strains, with Nes-cre displaying a the most phenotype hits (21) and Vil-cre displaying the least (2). Physiology phenotypes were most common, with both Nes-cre and Ins2-cre showing 19 and 13 abnormalities, respectively. As noted above, metabolic phenotypes in Nes-cre mice has been reported by others (Galichet et al. 2010; Giusti et al. 2014), and a recent paper suggests these phenotypes are due to presence of the hGH minigene (Declercq et al. 2015). The same hGH minigene is also found in the Ins2-cre allele, possibly explaining the metabolic phenotypes observed. This line is reported to develop age-related impaired glucose tolerance of unknown etiology(Lee et al. 2006), and thus the presence of the hGH minigene should be explored as a possible explanation. Both Alb-cre and Lck-cre also carry this minigene, but do not show the same number of phenotypic hits, suggesting that expression of the minigene varies between transgenes, and thus it cannot be assumed that its presence alone is necessarily confounding. It is interesting that Vav1-icre displayed only two hits despite landing in the *Commd10* gene, for which a KO allele is homozygous lethal. This is consistent, however, with the IMPC phenotyping data (www.mousephenotype.org), which shows no significant phenotypes in *Commd10^tm1a(EUCOMM)Wtsi/+^* mice. By contrast, the Wnt1-cre transgene inserts into the histone gene *H2afv* and shows 11 phenotypic hits which span several domains (Figure 5), including four significant behavioral phenotypes. Currently, there are no reports of targeted mutations of phenotypes for this gene.

**Figure 5.**
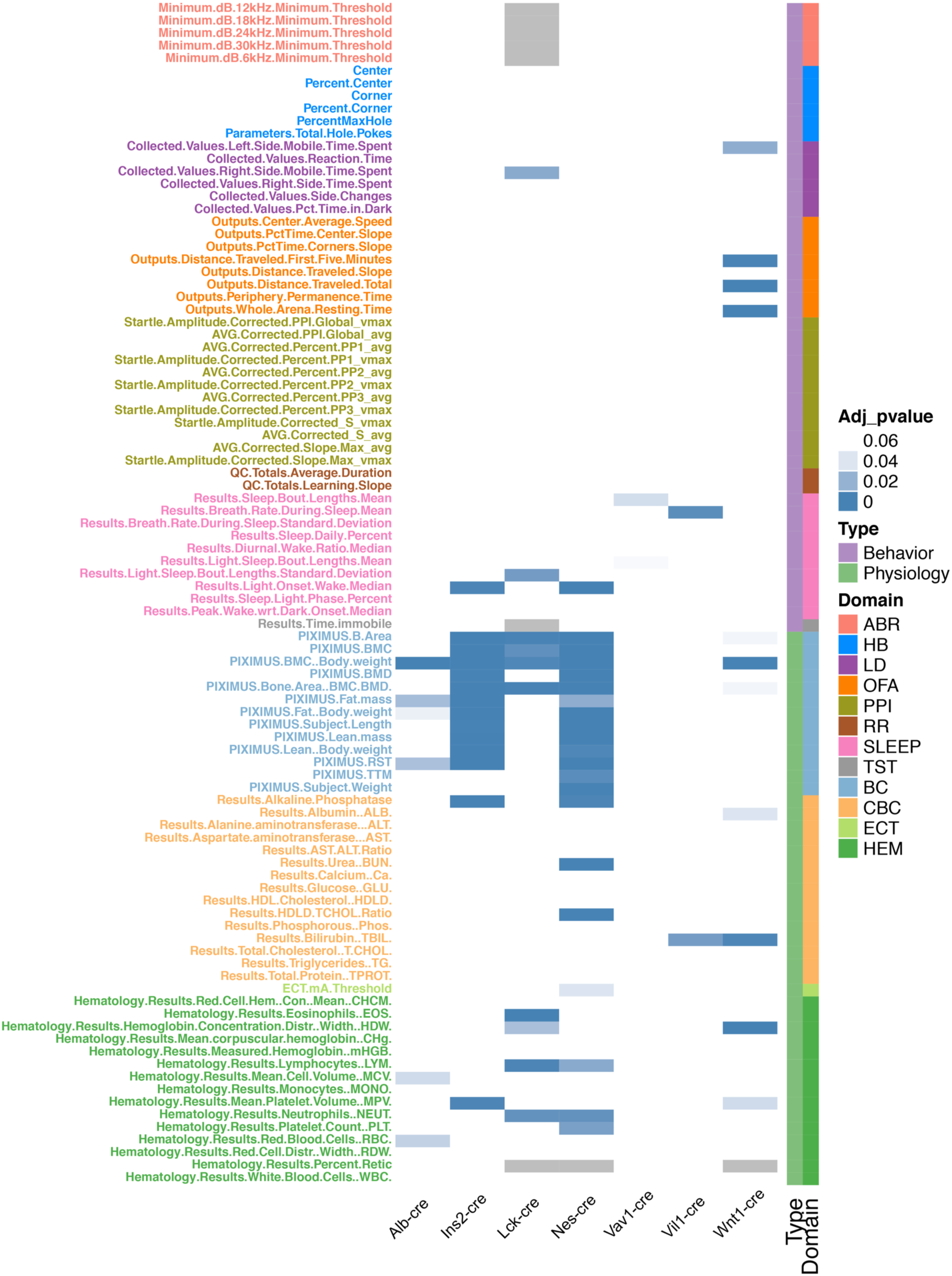
Physiology and behavioral testing of the cre transgenic lines in the KOMP pipeline. Mice were tested in 12 phenotypic domains spanning behavior and physiology (color coded, right bar). Each test is color coded (right bar) and grouped broadly into behavior (purple) and physiology domains (green). Significant differences from controls are shows in the heat map (FDR corrected p-values. Individual output parameters are listed and color coded on the left y-axis. Tests with no data are shown in gray. ABR: Auditory Brainstem Response; HB: Hole Board; LD: Light/Dark Transition; OFA: Open Field Assay; PPI: Pre-pulse inhibition; RR: Rotarod; SLEEP: Piezoelectric Sleep/Wake; TST: Tail Suspension Test; BC: Body Composition; CBC: Clinical Biochemistry; ECT: Electroconvulsive Seizure Threshold; HEM: Hematology.

## DISCUSSION

Despite widespread use of transgenic lines in the scientific community, the consequence of random transgene insertion in these lines is largely unknown. Here we show, expanding on prior work, that TLA represents a rapid and efficient means to precisely identify the site of insertion, and critically, the corresponding molecular consequences. These events are associated with structural variations, primarily deletions and duplications, including a deletion of greater than 1Mb and a complex structural variation that includes a simultaneous duplication, deletion, and inversion. While TLA simplifies the discovery process, reconstruction of the full transgene structure is difficult due to the high copy concatemerization of the transgene, coupled with complex structural variation that can accompany insertion. Given these contigs are built using relatively short read sequencing technology, including several instances where the fusion was covered by one read, it is critical to validate each putative fusion fragment using PCR-sequencing, and we indeed found small differences in the actual fusion sequence in a few lines. Targeted long-read sequencing approaches (PacBio, Oxford nanopore), might prove to be a useful complement to TLA for full characterization of transgenic alleles. However, such reconstruction is not necessary for typical use, as the key elements of chromosome location, break point, and structural variation are easily obtained with TLA and PCR validation alone.

In contrast to a recent similar screen on a small number of Cre driver lines(Cain-Hom et al. 2017), we found a surprisingly high percentage (50%) disrupt annotated genes, the majority of which are protein coding. Many of these genes, when mutated, are known to result in observable phenotypes, including four that are homozygous lethal. At face value, this is unexpected as only 3% of the genome is protein coding, and thus we would expect a similar hit rate with truly random insertion. However, many of these insertions are accompanied by substantial deletions, which would increase the odds of hitting a gene. In addition, our set of transgenic lines in not an unbiased selection of random transgenic animals; they are lines selected for robust transgene expression and activity. Genic regions of the genome are likely to support active transgene expression, as opposed to intergenic stretches and regions of heterochromatin. While we do observe that larger transgenes derived from BAC (or similar) constructs, which contain a larger complement of genetic elements required for proper expression, can insert into and/or delete coding genes (e.g. Cspg4-cre), our sample size is too small to determine if the rate is different than that of smaller transgenic constructs.

TLA-based discovery of transgenic insertion sites provides a number of practical benefits that should improve quality control for both public repositories and the end user. For example, allele-specific assays can be developed at the integration site to distinguish all genotype classes, allowing for homozygous mating strategies unless precluded by insertional mutagenesis. End users of Cre lines can use knowledge of the genetic locus before attempting to mate to a floxed target allele that is linked to the Cre line, selecting an alternative unlinked line, or scaling their breeding to assure identification of rare recombinants.

Our findings further illustrate the need to use proper controls in all experiments that include transgenic lines. Several studies have shown that expression of Cre itself can have phenotypic or toxic effects (Loonstra et al. 2001; Naiche and Papaioannou 2007; Bersell et al. 2013; Lexow et al. 2013). Given these findings it is clear that animals/embryos expressing Cre alone must be included as a control, and our results provide additional evidence that this control strategy is essential. Therefore, the potentially confounding impact of frequent insertional mutagenesis in Cre driver lines can be managed with the proper use of controls, depending on the research question and phenotype. For transgenic lines that are employed as disease models, a Tg-only control is not possible. In many cases, the original publication included results from multiple founders corroborating the findings of the line that ultimately became the “standard” for subsequent studies. It is interesting, however, that most studies published with an “established” model do not include the same level of independent corroboration, despite significant differences in study design, including analysis of additional phenotypes, inclusion of additional mutant alleles, and/or the use of a distinct genetic background. It is plausible that in those scenarios, effects of insertional mutagenesis not seen in the original publication might manifest, confounding the interpretation of the data. Typically, multiple alleles are not deposited in a public biorepository for distribution or retained at all, and thus reproduction of results with independent transgenic lines is impossible, notwithstanding the practical and financial challenge of reproducing every study with multiple transgenic lines. Thus, it seems prudent, and now feasible, for investigators to determine the insertion site of the transgenic line used in their study if independent corroboration is not possible.

One potential use of TLA is to confirm the content of a given transgenic line, providing a level of quality control not available through other means. This includes clarification of the specific details of the constructs components (e.g. hGH or MT1 minigenes) that are either omitted or reported incorrectly. It is worth noting that in the case of both “polyA” signals reported here, a careful review of the literature clearly shows that the original content of the vectors is correctly reported (Sauer and Henderson 1990; Orban et al. 1992), but this information was omitted or incorrectly cited in subsequent descriptions of the construct or mouse strain. Given the number of years and hands involved, this type of “information mutation” is not surprising. Indeed, we have seen that ~20% of all lines submitted to the JAX Repository carry alleles or have been bred to mouse strains not reported by the donating investigator. TLA provides an additional tool for assuring the content and nature of the allele for both investigators sharing their strain and for repositories distributing strains to scientists around the world.

Primary phenotyping of a subset of Cre drivers included in this study demonstrate the potential scope of “endogenous” phenotypes in transgenes in common use. While the impact of insertional mutagenesis is clear, for most KO alleles only homozygous mutants are carefully phenotyped, and thus potential confounding heterozygous phenotypes are unclear. Moreover, some transgenes delete multiple genes, and the combinatorial effect on phenotype would require independent evaluation. Finally, the transgene-specific caveats of passengers (minigenes, genes on BACs, *E.coli* genome, etc.) require specific testing. As noted above, the use of proper controls can mitigate most concerns arising from insertional mutagenesis, passenger cassettes, or transgene toxicity, assuming it does not directly impact the phenotype of interest. However, with the emergence of high-throughput phenotyping pipelines (Dickinson et al. 2016; Karp et al. 2017; Meehan et al. 2017), it is now feasible to broadly characterize the phenotypes of a larger collection transgenic tool lines, perhaps in parallel to insertion site discovery.

Given our findings, and the potential caveats implied for the use of transgenes, it is tempting to suggest that the community should move away from making, and ultimately using, lines generated by random transgenesis. For Cre lines, knockin alleles targeting the endogenous locus of a desired driver gene has the added potential advantage of providing greater specificity, desirable given the high rate of off-target activity seen in many transgenic lines (Heffner et al. 2012). To this end, the EUCOOMTOOLs program has produced hundreds of new Cre driver lines using this strategy (Murray et al. 2012; Rosen et al. 2015). However, this typically comes at the cost of haploinsufficiency at the driver locus, often a gene that is part of a pathway critical to the development of the cell type or tissue in question. Expression levels in a knockin might be lower than that of a multicopy transgene, thus sacrificing effectiveness for specificity. The use of neutral locus docking sites or targeted transgenesis facilitated by CRISPR can avoid the mutagenic risk associated with random insertion, but the former is typically a single copy event and the latter is relatively untested. Thus, while alternatives to random transgenesis exist, they come with their own caveats and do not necessarily provide a suitable alternative. Rather, given the impact of discoveries enabled by transgenic lines, knowledge of the transgenic insertion site is best viewed as one of many critical pieces of information that should be considered in an experimental design.

## METHODS

### Mice

All strains used for the TLA analysis were obtained from the Jackson Laboratory Repository, 4 of which are distributed from the JAX Mouse Mutant Research and Resource Center (MMRRC). The specific mouse strains and JAX Stock # (and MMRRC Stock # if applicable) are available in Supplemental Table 1. All procedures and protocols (see Phenotyping below) were approved by The Jackson Laboratory Animal Care and Use Committee, and were conducted in compliance with the National Institutes of Health Guideline for Care and Use of Laboratory Animals.

### Isolation of splenocytes

Splenocytes were isolated from each line as previously described (de Vree et al. 2014). In brief, the spleens were dissected and stored on ice. A single cell suspension was made using a 40 micron mesh filter suspending the cells in 10% Fetal Calf Serum (FCS)/Phosphate Buffered Saline (PBS). Following centrifugation at 4°C at 500 × g for 5 min. The supernatant was discarded, the pellet dissolved in 1ml 1 × Pharm Lyse (BD Biosciences) and incubated at room temperature for 3 min to lyse splenic erythrocytes. To terminate the lysis reaction, 0.5 ml phosphate buffered saline (PBS) was added followed by centrifugation at 4°C, 500 × g for 5 min. The supernatant was discarded and the pellet resuspended in 0.5 ml PBS. After one final centrifugation step for 2 min, the supernatant was discarded and cell pellet resuspended in 1 ml freeze medium (PBS with 10% Dimethyl Sulfoxide and 10%fetal calf serum). The samples were stored at minus 80°C until shipment for TLA processing.

### TLA procedure

Targeted locus amplification (TLA) was performed as previously described. (de Vree et al. 2014) In brief, Cells were crosslinked using formaldehyde, after which the DNA was digested using the restriction enzyme NlaIII (CATG). Subsequently the sample was ligated, crosslinks were reversed and the DNA was purified. To obtain circular chimeric DNA molecules for PCR amplification, the DNA molecules were trimmed with NspI and ligated at a DNA concentration of 5 ng/μl to promote intramolecular ligation. NspI has a RCATGY recognition sequence that encompasses the CATG recognition sequence of NlaIII, which ensures only a subset of NlaIII (CATG) sites were (re-) digested, generating DNA fragments of approximately 2 kb and allowing the amplification of entire restriction fragments. After ligation the DNA was purified, and eight 25-μl PCR reactions, each containing 100 ng template, were pooled for sequencing. Sequences of the inverse primers, which were designed using Primer3 software38, can be found in Supplementary Table 4.

### Mapping and sequence alignment

The primer sets were used in individual TLA amplifications. PCR products were purified and library prepped using the Illumina NexteraXT protocol and sequenced on an Illumina Miseq sequencer. Reads were mapped using BWA-SW, which is a Smith-Waterman alignment tool. This allows partial mapping, which is optimally suited for identifying break-spanning reads.The mouse mm10, rat rn5, cow bosTau8, SV40 GCF_000837645.1, rabbit oryCun2, chicken Galgal4 and human genome version hg19 were used for mapping. Identified TG sequences based on these alignments have been confirmed in this paper.

### Sequence validation

Breakpoint-spanning reads-All TG integration fusion sites were confirmed by PCR amplification and sequence analysis. The extended reads were analyzed for GC content using ENDMEMO Software (http://www.endmemo.com/bio/gc.php) and PCR primers were designed using Primer 3 software (Untergasser et al. 2012) to optimize for size and GC content. If the fusion product was larger than 900 bp the fusion site was confirmed using at least 2 sets of primers for the long read as well as an internal read to insure adequate coverage of the integration site. PCR amplicons with suitable products were purified and Sanger sequenced.

### QPCR analysis

Genomic DNA isolated from tail biopsies were used to analyze loss of allele (LOA) or relative concentration qPCR on Applied Biosystem’s ViiA 7 (Applied Biosystems, Foster City, CA USA). LOA assay design: The premise behind the LOA assay assumes a one-copy difference between a Transgene Insertion and the wild-type (WT) sample, while a gain of allele can be used to show a duplication of the genomic target region. Based on transgene integration site sequences and resultant deletions or duplications, target genomic region q-PCR 5’ nuclease assays were designed using PrimerQuest software (Integrated DNA Technologies). The internal reference control *Apob* probe contains a VIC (4,7,2′-trichloro-7′-phenyl-6-carboxyfluorescein) reporter dye, (ABI, Applied Biosystems, Foster City, CA USA) while all experimental assays use FAM (6-carboxyfluorescein) labeled probes for detection. An NFQ-MGB (*Apob*) dark or Zen/Iowa Black FQ quencher (IDT) is used for all assays. Primer and probe sequences are provided in Supplementary Table S3.

QPCR samples were then analyzed in triplicate and Cq values for the samples and the internal reference (*Apob*) were calculated using Viia7 Software (QuantStudio™ Software V1.3, ABI, Applied Biosystems, Foster City, CA USA). The means of the Cq values were used to calculate ΔCq values and these were then used to calculate relative copy number of the recombinant region using the 2–ΔΔCq formula (Livak and Schmittgen 2001).

### Phenotyping

We employed a modified version of the IMPReSS pipeline (www.mousephenotype.org/impress) for high-throughput clinical phenotyping assessment, which was developed under the IMPC program (Dickinson et al. 2016; Karp et al. 2017; Meehan et al. 2017), de Angelis, Nat Genet. 2015 September; 47(9): 969–978. doi:10.1038/ng.33). The following seven lines (and genotypes) were characterized: Alb-cre (HOM), Ins2-cre (HOM), Lck-cre (HOM), Nes-cre (HEMI), Vav1-icre (HEMI), Vil-cre (HEMI) and Wnt1-cre (HEMI). Control mice are from a pool consisting of C57BL/6J WT mice and non-carrier (NCAR) controls from the colony for lines maintained in a HEMI x NCAR breeding scheme (Wnt1-cre, Vav1-icre, Vil-cre, and Nes-cre). For each mouse strain, 8 male and 8 female transgenic animals, NCAR controls, or C57BL/6J mice were processed through the JAX Adult Phenotyping Pipeline. Full details of the JAX Adult Phenotyping Pipeline can be found on the IMPC website (www.mousephenotype.org/impress/procedures/12). Briefly, mice were received into the pipeline at 4 weeks of age, body weight was collected weekly, and assays were performed weekly from 8 to 18 weeks of age, ordered such that the least invasive, behavioral testing was performed first. The specific assays and age in weeks that the assay was performed in this study are as follows:

Open Field (OFA) (8 wks)
Light-Dark Transition (LD), Holeboard (HB) (9 wks)
Acoustic Startle/Pre-pulse Inhibition (PPI) (10 wks)
Tail Suspension, Electrocardiogram, Rotarod (11 wks)
Body Composition (BC), (14 wks)
Piezoelectric Sleep/Wake (SLEEP) (15 wks)
Auditory Brainstem Response (ABR) (4M + 4F) (16 wks)
Electroconvulsive Seizure Threshold (ECT) (17 wks)
Terminal collection including Hematology (HEM), Clinical Biochemistry (CBC)

### JAX-specific sleep test

Sleep and wake states were determined using the PiezoSleep System (Flores et al. 2007; Donohue et al. 2008; Mang et al. 2014). The system is comprised of plexiglass cages lined with piezoelectric films across the cage floor that detect pressure variations. Signal features sensitive to the differences between the sleep and wake states are extracted from 8-second pressure signal segments, and classification is automatically performed every 2 seconds using overlapping windows. From this, the following parameters are calculated: Sleep bout lengths (light phase, dark phase, 24 hour mean), breathing rate, breathing rate during sleep, percentage daily sleep (light and dark phase), and diurnal wake ratio.

### Statistical analysis

Linear mixed models (LMM) were performed to identify phenotypic associations from high throughput phenotyping experiments. Sex, weight and batch were a significant source of variation for continuous phenotypes. In the linear mixed model, explanatory factors including sex, weight and mutant genotype were treated as fixed effects, while batch (date of test) was treated as a random effect adding variation to the data (see Equation 1).

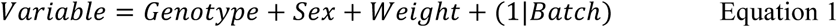

Parameters from the mixed model are estimated using the method of restricted maximum likelihood (REML). Adjusted p-values were calculated from nominal p-values in mixed models to control for False Discovery Rate (FDR). All data analysis was performed using R.

## DATA ACCESS

All sequence date will be submitted to the NCBI Sequence Read Archive (SRA; http://www.ncbi.nlm.nih.gov/sra) Phenotypic data have been submitted to the Mouse Phenome Database at phenome.jax.org

## ACKNOWLEDGEMENTS

This work was supported in part by the Office of the Director, National Institutes of Health and the National Human Genome Research Institute of the National Institutes of Health under award numbers R24OD011 190, U42OD011185, UM1OD023222, and HG006332. The authors thank Brianna Caddle and Larry Bechtel for their technical assistance in isolating spleen cells, and Kevin Peterson for his helpful and thoughtful comments on the manuscript.

## SUPPLEMENTARY INFORMATION

**Supplementary Table 1.** Detailed summary of results for each of the 40 lines included in this study

**Supplementary Table 2.** Primers and expected PCR product sizes used for validation of insertion sites.

**Supplementary Table 3**. qPCR probes used for LOA validation.

**Supplementary Table 4**. Primers used for TLA amplifications.

**Supplementary Table 5.** Table of statistical results for mouse phenotyping.

